# Genome-wide detection of somatic mosaicism at short tandem repeats

**DOI:** 10.1101/2023.11.22.568371

**Authors:** Aarushi Sehgal, Helyaneh Ziaei-Jam, Andrew Shen, Melissa Gymrek

## Abstract

**Motivation:** Somatic mosaicism, in which a mutation occurs post-zygotically, has been implicated in several developmental disorders, cancers, and other diseases. Short tandem repeats (STRs) consist of repeated sequences of 1-6bp and comprise more than 1 million loci in the human genome. Somatic mosaicism at STRs is known to play a key role in the pathogenicity of loci implicated in repeat expansion disorders and is highly prevalent in cancers exhibiting microsatellite instability. While a variety of tools have been developed to genotype germline variation at STRs, a method for systematically identifying mosaic STRs (mSTRs) is lacking.

**Results:** We introduce prancSTR, a novel method for detecting mSTRs from individual high-throughput sequencing datasets. Unlike many existing mosaicism detection methods for other variant types, prancSTR does not require a matched control sample as input. We show that prancSTR accurately identifies mSTRs in simulated data and demonstrate its feasibility by identifying candidate mSTRs in whole genome sequencing (WGS) data derived from lymphoblastoid cell lines for individuals sequenced by the 1000 Genomes Project. Our analysis identified an average of 76 and 577 non-homopolymer and homopolymer mSTRs respectively per cell line as well as multiple cell lines with outlier mSTR counts more than 6 times the population average, suggesting a subset of cell lines have particularly high STR instability rates.

**Availability:** prancSTR is freely available at https://github.com/gymrek-lab/trtools.

**Documentation:** Detailed documentation is available at https://trtools.readthedocs.io/

## Introduction

Population-level heterogeneity generally arises due to germline mutations that occur before the formation of the zygote and are inherited by all cells in the offspring. However, heterogeneity within an individual may also exist due to somatic mutations that occur post-zygotically in only a sub-population of cells (reviewed in (Youssoufian and Pyeritz, 2002)). Somatic mosaicism has long been known to play a key role in cancer (reviewed in (Stratton *et al*., 2009)), and has also been implicated in a range of non-neoplastic disorders (e.g., Proteus Syndrome (Cohen, 1993), Neurofibromatosis Type 1 (Ruggieri and Huson, 2001) and CLOVES syndrome (Kurek *et al*., 2012)). Somatic mosaicism is also a hallmark of conditions resulting in DNA repair deficiencies, such as Xeroderma Pigmentosum (Cleaver, 1969). Beyond its role in disease, accumulation of somatic mutations is likely a widespread phenomenon occurring in healthy individuals across the course of their lifetime (Fernández *et al*., 2016).

High-throughput sequencing offers the potential to perform genome-wide detection of somatic mosaicism, but also presents important technical challenges (Dou *et al*., 2018). To distinguish somatic mutations from germline variants or technical artifacts, a matched control sample is often required to serve as a baseline. Further, in cases where the somatic mutation is present in a small fraction of cells, ultra high coverage data is needed to detect the event (Breuss *et al*., 2022). A variety of methods have been developed to address these challenges (e.g. MrMosaic (King *et al*., 2017), MosaicForecast (Dou *et al*., 2020), and DeepMosaic (Yang *et al*., 2023)). These methods leverage allele fractions, read-based phasing, and other read-level features to accurately distinguish true mosaic variants. However, existing methods in some cases still require matched control samples and focus largely on detecting mosaic single nucleotide polymorphisms (SNPs) or in some cases mosaic copy number variants (e.g. Montage (Glessner *et al*., 2021)).

Short tandem repeats (STRs), consisting of 1-6bp sequences repeated in tandem, comprise more than 3% of the the human genome (Lander *et al*., 2001), and exhibit rapid germline mutation rates (Sun *et al*., 2012). Somatic instability of STRs, also known as microsatellite instability (MSI), is a hallmark of certain cancers such as Lynch Syndrome (reviewed in (Lynch *et al*., 2009)). Recent work suggests mutation rates of approximately 10^*−*4^-10^*−*3^ mutations per STR in non-MSI cancers, with rates more than in the case of MSI (Fujimoto *et al*., 2020). Additionally, somatic mutation of STRs in the brain has been implicated as a key driver of pathogenicity in some repeat expansion disorders including Huntington’s Disease (Swami *et al*., 2009).

Detection of somatic mosaicism at STRs from sequencing data is particularly challenging, as these regions may exhibit high error rates due to PCR artifacts (Raz *et al*., 2019) making it difficult to distinguish true somatic mutations from errors. STR-specific genotyping methods have been developed for germline genotyping that address this challenge (e.g. HipSTR (Willems *et al*., 2017) and ExpansionHunter (Dolzhenko *et al*., 2017)), but these are not designed to detect somatic events. Previous studies performed genome-wide analysis of somatic STR instability in the context of cancer (Hause *et al*., 2016; Kim et al., 2013; Fujimoto et al., 2020), but relied on comparing sequencing from tumors with matched normal samples. Further, somatic events were detected either using custom analysis pipelines not packaged as a separate tool (Kim *et al*., 2013) or were based on heuristics rather than hypothesis testing frameworks (Salipante *et al*., 2014).

Here, we introduce prancSTR, a novel method for detecting mosaic STRs (mSTRs) from next-generation sequencing data without the need for a matched control sample. prancSTR models observed reads as a mixture distribution and infers the maximum likelihood mosaic fraction and the copy number of the mosaic vs. germline alleles. We show that prancSTR accurately identifies mSTRs in simulated data and validate mSTRs inferred from short reads with orthogonal long read data. Finally, we apply prancSTR to 460 whole genome sequencing (WGS) datasets from the 1000 Genomes Project derived from lymphoblastoid cell lines (LCLs) to characterize genome-wide mSTRs in different populations. Overall, prancSTR provides a robust method to identify mSTRs from existing high throughput sequencing datasets.

## Methods

### prancSTR overview

#### Baseline model

prancSTR is designed to identify mSTRs at one locus at a time. It takes as input STR genotypes and metadata computed by an existing genotyper and outputs candidate mSTRs (**Fig. 1A**). While designed to work downstream of HipSTR (Willems *et al*., 2017), prancSTR can theoretically process output from any STR genotyping tool as long as it returns estimated diploid repeat lengths and the observed distribution of copy numbers across all reads aligning to a locus.

**Fig. 1.**
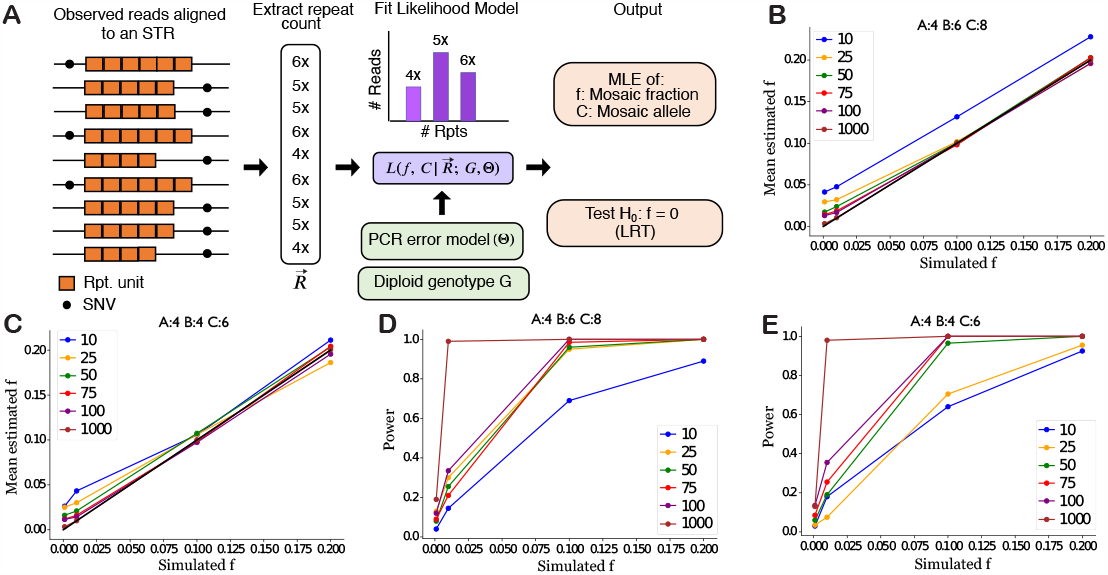
prancSTR overview and validation. (A) Overview of the prancSTR method. The copy numbers observed in each read aligned to a target STR are extracted to a vector 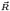, from which prancSTR obtains maximum likelihood estimates for the mosaic allele (*C*) and mosaic allele fraction (*f*), and a P-value testing *H*_0_ : *f* = 0. (B-C) Simulated vs. estimated values of *f* . We simulated mSTRs under a range of coverage levels and values for *f* for cases in which the germline genotype is heterozygous (B) or homozygous (C). Dots represent the mean estimated *f* value from 200 simulations. The black line denotes the x=y diagonal. (D-E) Power to detect mSTRs. Power is computed as the percent of simulations for which *P <* 0.05. For B-E, lines denote different coverage levels, where coverage gives the total number of reads spanning the STR of interest. Simulated values for *A, B*, and *C* are denoted at the top of each panel. Panels here are based on simulated read vectors 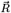. Similar results for simulations based on raw reads are shown in Supplementary Figs. 3-4.

At each STR locus, prancSTR takes as input a vector of the observed repeat copy number in each read, 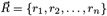, where *r*_*i*_ is the number of copies of the repeat observed in the *i*th read. For each locus, let *⟨A, B⟩* denote the diploid germline genotype, where *A* and *B* give the copy number of the repeat unit on each allele. Let *f* denote the fraction of chromosome copies harboring an additional allele *C* resulting from a mosaic mutation, and Θ represent additional error parameters described below. If the somatic mutation occurred on the haplotype containing allele *B*, we would expect 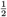 of chromosome copies to contain allele *A*, 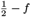 to contain allele *B*, and *f* to contain allele *C*. Assuming each observed read is independent, we can then write the following likelihood equation:

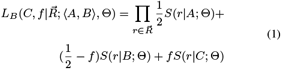

where *L*_*B*_ denotes the likelihood of *C* and *f* in the case that the mosaic allele occurred on the haplotype with allele *B. S*(*r*|*G*; Θ)gives the probability to observe *r* copies of the repeat in a read given it originated from an allele with *G* copies assuming stutter error model Θ. This term is computed based on the error model used in HipSTR (Willems *et al*., 2017):

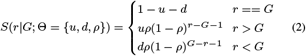

where *u* and *d* give the probability for a read to contain a stutter error resulting in a repeat expansion or contraction, respectively, and error step sizes are assumed to follow a geometric distribution with parameter *ρ*. We assume here that *u, d*, and *ρ* are known for each locus as these can be estimated from existing data using other methods (Willems *et al*., 2017; Kristmundsdottir *et al*., 2020).

In practice with short reads we are unable to determine the haplotype of origin (either *A* or *B*) of the mosaic allele. Therefore below we aim to identify *C* and *f* that maximize the log likelihood over two possible cases:

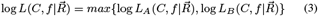

#### Likelihood maximization and hypothesis testing

The goal of prancSTR is to find values for *C* and *f* that maximize Equation 3. We assume the underlying stutter model Θ and diploid genotype *⟨A, B⟩* are known and can be obtained from HipSTR’s output. We then use an iterative algorithm to estimate *C* and *f* :

1. Initialize the value of *f* to 0.01.
2. Compute the log-likelihood for each possible value of *C*, given *f* from step 1. We restrict our search for *C* to (min 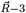*−*3, max 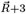 +3). Return the value of *C* that maximizes the log-likelihood.
3. Find the value of *f* that maximizes the log-likelihood given *C* from step 2. This step is performed using Sequential Least Squares Programming (SLSQP) (Kraft, 1988) restricting *f* to be between 0 and 0.5.
4. Repeat until convergence.

In practice, the read vector 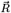 is obtained from the MALLREADS format field from HipSTR VCF files. We exclude STR calls from analysis if: they have coverage of 0, have missing genotypes, have 0 read support in MALLREADS for the called diploid genotype, or if there is only evidence in MALLREADS of reads from a single allele.

After obtaining the maximum likelihood estimates 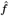 and *Ĉ*, prancSTR tests the null hypothesis *H*_0_ : *f* = 0 (no mosaicism) at each STR genotyped in each sample. We compute the likelihood ratio test statistic *λ*_*LR*_:

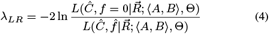

Finally, we use the fact that *λ*_*LR*_ *∼ χ*^2^(2) to obtain a P-value testing *H*_0_ : *f* = 0 at each STR in each sample. Here, since the null hypothesis (*f* = 0) falls on the boundary of the parameter space, we use a null consisting of a mixture distribution of a point mass at 0, with 50% probability, and a chi-square distribution with 2 degrees of freedom, also with 50% probability (Nielsen and Wakeley, 2001).

#### Simulating vectors of observed repeat counts

In our first simulation strategy, we simulated vectors of observed repeat counts for a single locus according to the baseline model described above under various parameter settings. The resulting read count vectors, as well as the known values of *A, B*, and Θ, were used as input to prancSTR’s likelihood estimation procedure. In all cases, the mosaic allele fraction *f* was set to one of [0.001, 0.01, 0.1, 0.2], the total number of reads *N* was set to one of [10, 25, 50, 75, 100, 1000], and stutter parameters were fixed at *ρ* = 0.9, *u* = 0.02, *d* = 0.02. We tested settings in which the true diploid genotype was set either to *⟨*4, 4*⟩* (homozygous) or *⟨*4, 6*⟩* (heterozygous). The value of the mosaic allele *C* ranged from 6-10 repeats.

For each tested setting, we performed 200 simulations. Power was estimated as the percentage of simulation rounds for which prancSTR returned a significant P-value (P<0.05). Notably this captures relative power differences across settings but is not reflective of the absolute power in genome-wide analyses, in which a more stringent P-value threshold is required to account for multiple hypothesis testing. To evaluate false positive rates, we performed simulations with *f* set to 0 and similarly returned the percentage of simulation rounds with significant P-values.

#### A method for simulating error-prone next-generation sequencing reads at STRs

For our second simulation strategy, we developed a novel simulation framework, simTR, which simulates raw sequencing reads according to a specified coverage level and error model using user-defined repeat alleles. simTR is a wrapper built around ART (Huang *et al*., 2012), an existing, open source, next generation sequencing read simulator. ART creates simulated reads that account for generic insertion and deletion mutations. However, stutter errors (additions or deletions of one or more repeat units introduced during PCR) characteristic of STRs are not specifically modeled. simTR adds to ART by incorporating stutter errors into the simulated reads, in addition to existing indel mutations. Stutter errors are incorporated based on the HipSTR error model described in Equation 2.

simTR takes as input a genome file (fasta format), the genomic coordinates of the target STR, and stutter parameters (*u, d*, and *ρ*). Users may also specify optional parameters to set the desired coverage, to generate paired-end vs. single-end reads, the mean and standard deviation of the sequencing fragment lengths, and the window size around the STR from which to simulate reads. It creates intermediate fasta files with separate entries to represent the different possible observed repeat lengths that could result from PCR stutter. It then invokes ART to simulate reads from the different fasta entries at rates proportional the expected proportion of each allele based on the input stutter parameters. Finally, it outputs simulated reads in fastq format which can be used for benchmarking downstream tools.

To evaluate the entire prancSTR pipeline starting from raw reads, we applied simTR to simulate reads at a target set of mSTRs under a range of settings. Simulated reads were aligned to a reference genome (hg38) using BWA MEM (Li, 2013) version 0.7.12-r1039. The resulting reads were used as input to HipSTR v0.6.1 for genotyping the target STRs using non-default options min-reads 5 and stutter-in to provide a file with simulated stutter error parameters. The VCF output by HipSTR was then used as input to prancSTR to estimate *C* and *f* .

We tested this pipeline on two example STR loci: (1) CSF1PO, a tetranucleotide (ATCT)_*n*_ CODIS marker annotated by the National Institute of Standards and Technology (NIST) (https://strbase-archive.nist.gov/str_CSF1PO.htm); (hg38 chr5:150071324-150081375), and (2) a (CGG)_*n*_ repeat in *CBL* (hg38 chr11:119206289-119206322). For CSF1PO, the true diploid genotype was set to either *⟨*13, 11*⟩* (heterozygous) or *⟨*13, 13*⟩* (homozygous), and the mosaic allele was set to 15. For the *CBL* repeat, the true diploid genotype was set to either *⟨*11, 14*⟩* or *⟨*11, 11*⟩*, and the mosaic allele was set to either 9 or 10. We tested a range of values for *N* (10, 25, 50, 75, 100, 1000) and *f* (0, 0.01, 0.1, 0.2) and set stutter parameters *ρ* = 0.9, *u* = *d* = 0.02 to simulate 150bp paired-end reads. Example IGV (Thorvaldsdóttir *et al*., 2013) screenshots for simulated reads at mSTRs are shown in **Supplementary Fig. 1**.

#### Obtaining estimated stutter parameters from real WGS datasets

We previously performed genome-wide STR genotyping using HipSTR on high-coverage PCR-free WGS for 3,202 individuals from the 1000 Genomes Project and 348 PCR+ samples from the H3Africa cohort (Ziaei Jam *et al*., 2023). Per-locus stutter parameters estimated by HipSTR were extracted from VCF files (INFO fields INFRAME_UP, INFRAME_DOWN, and INFRAME_PGEOM) for individuals from the Yoruban population (1000Genomes) and H3Africa cohorts separately using bcftools (Danecek *et al*., 2021) v1.10.2.

#### Implementation

prancSTR and simTR are implemented in Python as an open source command line tool and are available as part of the TRTools (Mousavi *et al*., 2021) package.

#### Characterizing mSTRs in the 1000 Genomes Project

We applied prancSTR to identify candidate mSTRs in 1000 Genomes Project samples based on previously obtained HipSTR calls (Ziaei Jam *et al*., 2023). These calls had already been filtered to exclude loci with call rate less than 75%, loci with genotypes not matching Hardy-Weinberg expectation (p*<*1e-06), and loci overlapping segmental duplications in the human genome. prancSTR output was filtered to include candidate mSTRs with: at least three reads supporting the identified mosaic allele *C*, read depth at least 10, mSTRs with *f ≤* 0.3 (larger *f* indicates likely heterozygous sites), HipSTR quality score *≥* 0.8. To adjust for multiple hypothesis correction (one test per locus), we applied the Benjamini-Hochberg (Benjamini and Hochberg, 1995) method to identify mSTRs at a false discovery rate of 5%.

Samples with outlier numbers of mSTRs were identified as those with mSTR counts more than two standard deviations above the mean across all individuals in each population. WGS sequencing coverage and EBV coverage for each sample was obtained from the 1000 Genomes Project website: http://ftp.1000genomes.ebi.ac.uk/vol1/ftp/data_collections/1000G_2504_high_coverage/1000G_2504_high_coverage.sequence.index and ftp://ftp.1000genomes.ebi.ac.uk/vol1/ftp/technical/working/20130606_sample_info/20130606_sample_info.txt.

#### Validating mSTRs from NA12878 using PacBio HiFi long reads

Aligned reads (BAM) for NA12878 based on PacBio HiFi long reads were obtained from Genome In A Bottle (https://ftp-trace.ncbi.nlm.nih.gov/ReferenceSamples/giab/data/NA12878/PacBio_SequelII_CCS_11kb/HG001_GRCh38/). We used the haplotag (HP) tag to partition the BAM into separate files containing reads for each haplotype. We then used HipSTR v0.7 to separately genotype reads from each haplotype using the following non-default parameters to enable running HipSTR on long reads: def-stutter-model, max-str-len 1000, max-flank-indel 1, use-unpaired, no-rmdup, min-reads 5, output-filters. We extracted the MALLREADS field from the HipSTR VCF file to examine support for each allele in PacBio reads for each haplotype. Analysis was restricted to loci with at least 10 spanning PacBio reads from each of the two haplotypes.

For comparison, we performed a similar analysis on all CEU samples, including NA12878, to assess how often PacBio reads (from NA12878) would appear to validate an mSTR that was identified in a different sample. For this analysis, we further filtered: (1) mSTRs where the mosaic allele was found on more than 80% of PacBio reads from a single haplotype (indicating the mosaic allele for a sample was most likely the NA12878 germline allele) and (2) unique STR loci identified as mSTRs in more than 5 samples, indicating the locus is potentially problematic. Finally, we excluded samples from the comparison if fewer than five mSTRs sites remained in a particular category, as the metrics computed are unreliable on low count numbers.

## Results

### Benchmarking prancSTR using simulated data

To evaluate prancSTR, we performed simulations using two strategies (**Methods**). First, to evaluate our likelihood maximization procedure, we simulated vectors of observed repeat counts in each read aligned to a locus 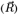 according to the baseline model described in **Methods**. In this case, we assumed the germline (diploid) genotype *⟨A, B⟩* is known, and use the ground truth values of *A* and *B* as well as the simulated read vectors as input to the maximum likelihood estimation of mosaic allele (*C*) and mosaic fraction (*f*). Second, to evaluate our end to end pipeline starting from raw reads, we used simTR to simulate reads for mSTRs under a range of conditions, which were used as input to HipSTR to infer the germline genotype and compute read vectors. HipSTR results were used as input for mosaicism detection.

We first evaluated prancSTR under the null setting of *f* = 0 to determine how often we falsely detect a significant mSTR. P-values returned by prancSTR are well-calibrated, following the expected uniform distribution in this case (**Supplementary Fig. 2A-B**). As expected, at a P-value threshold of 0.05, prancSTR falsely identifies approximately 5% of null simulation rounds as significant mSTRs (**Supplementary Fig. 2C-D**).

Next, we simulated mSTRs under a range of values for coverage and mosaic allele fraction and for cases in which the germline genotype is either homozygous or heterozygous. Using both simulation strategies, estimated values of the mosaic allele fraction 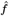 are highly consistent with simulated values (**Fig. 1B-C, Supplementary Figs. 3-4**). In cases that are underpowered (*f <* 0.02 and/or coverage 10*×*), prancSTR tends to slightly but consistently overestimate the mosaic allele fraction. In practice, these cases are unlikely to reach genome-wide significance.

As expected, power to detect mSTRs increases as a function of *f* and sequencing coverage in all simulation settings (**Fig. 1D-E, Supplementary Figs. 3-4**) with near perfect power at P<0.05 to detect mSTRs with *f >* 0.1 at loci with at least 50*×* coverage. In both simulation strategies, power is higher when the germline genotype is heterozygous vs. homozygous. This difference is more pronounced in results based on simTR simulations. In that case, this bias is partially explained by genotyping errors. We observed that cases where the simulated germline genotype is homozygous but the mosaic fraction is high are consistently misidentified by HipSTR as heterozygous sites, and therefore cannot be identified by prancSTR as mSTRs. We additionally evaluated the impact of the mosaic allele size on power. We observed that power increases with the absolute difference in length of the mosaic allele compared to the nearest germline allele (**Supplementary Fig. 5**). This is expected, since larger differences in size make it easier to distinguish true mosaic alleles from errors.

Finally, we evaluated the impact of sequencing errors at STRs on the ability to detect mSTRs from simulated read vectors under varying stutter model parameters meant to capture typical error rates in PCR+ (*∼*10% of reads) vs. PCR-free (*∼*1% of reads) data (**Supplementary Figs. 6-8**). As expected, with high stutter error rate, power is reduced in cases of low coverage and low mosaic fraction, and estimates of *C* and *f* show greater variability. This suggests mSTR detection will perform poorly on PCR+ short read data, where stutter error rates may often exceed expected mosaic fractions.

### Population-wide characterization of mSTRs

We next applied prancSTR to characterize population-wide trends of STR mosaicism. We focused on individuals from 1000 Genomes samples from the CEU (Northern Europeans from Utah; n=179), YRI (Yorubans from Nigeria; n=178), and CHB (Han Chinese; n=103) populations for which high-coverage PCR-free WGS is available (Byrska-Bishop *et al*., 2022). Notably, since the data is LCL-derived, identified mSTRs likely consist of a combination of true somatic mutations that existed before sample collection as well as mutations that have accumulated during cell line passages. After filtering (**Methods**), we identified an average of 76 (577) non-homopolymer (homopolymer) mSTRs per cell line (**Fig. 2A-B**). Overall, homopolymer mSTRs far outnumber non-homopolymers, and the majority of non-homopolymer mSTRs identified occur at loci for which the germline genotype is heterozygous (**Supplementary Fig. 9**). This trend is consistent across all populations analyzed.

**Fig. 2.**
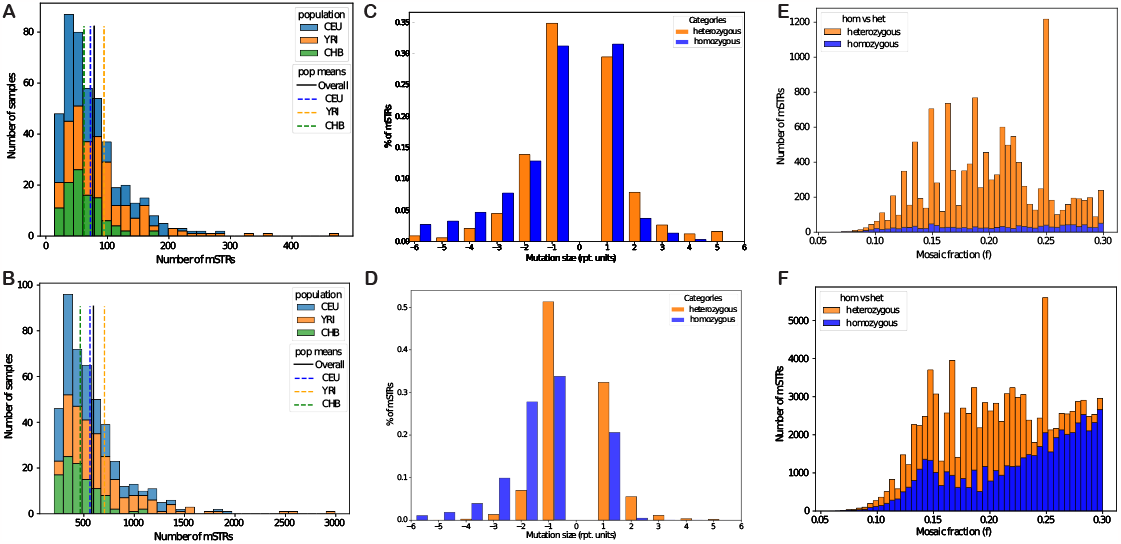
mSTR trends across populations. (A-B) Distribution of the number of mSTRs across different populations. The x-axis gives the number of mSTRs for a given population and the y-axis gives the number of samples. Data is shown for non-homopolymers (A) and homopolymers (B). Dashed colored lines (CEU=blue, YRI=orange, CHB=green) give population-specific means, and the black line denotes the overall mean. (C-D) Distribution of mSTR mutation sizes. The x-axis represents the mutation size, computed as the difference between the mosaic allele length and the closest germline allele. Positive mutation sizes indicate insertions and negative sizes indicate deletions. Data is shown for non-homopolymers (C) and homopolymers (D) and is for CEU only. Other populations showed similar trends. Blue represents homozygous loci and orange represents heterozygous loci. (E-F) Distribution of mosaic allele fraction (*f*) across mSTRs. Data is shown for non-homopolymers (E) and homopolymers (F) and is for CEU only. Bars are colored to denote the number of mSTRs occurring at homozygous (blue) vs. heterozygous (orange) sites.

We noticed substantial variation in mSTR counts across cell lines. The number of homopolymer and non-homopolymer mSTRs per cell line are highly correlated (**Supplementary Fig. 10**), with the correlation strongest when considering mSTR calls at germline heterozygous sites (Pearson r=0.97, two-sided P=9.6e-110 in CEU). We also identified 13, 8, and 4 cell lines from CEU, YRI, and CHB respectively with outlier mSTR counts (**Methods**) for both homopolymer and non-homopolymer mSTRs. Overall, these results suggest certain cell lines have higher rates of STR instability, potentially due to genetic or environmental factors. Variation in mSTR counts across cell lines is not significantly correlated with the number of sites considered or EBV virus count (two-sided P*≥* 0.05), and is only modestly correlated with sequencing coverage (Pearson r=0.21, two-sided P=0.039 for non-homopolymers and r=0.17, P=0.10; **Supplementary Fig. 11**). Passage numbers for these cell lines were not available at the time of writing, and so the impact of cell culture history, which is likely to play a role in mutation counts, could not be assessed.

We next investigated the distribution of the sizes of mosaic STR mutations. The majority of events (60.2% and 68.7% for non-homopolymer and homopolymer mSTRs respectively) result in insertions or deletions of a single repeat unit (**Fig. 2C-D**), although larger step sizes were observed. Mutation sizes are larger on average for mutations at STRs with homozygous vs. heterozygous germline genotypes and show an overall bias toward deletions vs. contractions. A similar deletion bias has been observed for somatic mutations at STRs in cancer (Fujimoto *et al*., 2020). However, both biases described above are far more pronounced at homopolymer loci, suggesting they my arise in part from erroneous mSTR calls (**Supplementary Fig. 12**). Indeed, inferred stutter error rates suggest deletion errors are more common than insertions (**Supplementary Figs. 7-8**), and large mutation step sizes at homozygous sites may reflect true heterozygous sites that were incorrectly genotyped.

We then examined the distribution of variant allele fractions (*f*) for detected mSTRs (**Fig. 2E-F, Supplementary Fig. 13**). In all cases, *f* distributions show peaks around 0.15-0.20, consistent with the range where we expect to have sufficient power (**Fig. 1D-E**), whereas true mosaic sites with higher *f* values are likely to be indistinguishable from heterozygous sites. Further, homopolymer mSTRs with high *f* values nearly all occur at homozygous sites, and the observed deletion bias is strongest overall for sites with high *f* values, indicating mSTRs with *f >* 0.2 may be enriched for false positive calls.

### Validating mSTRs in a deeply sequenced human sample

To further evaluate whether our pipeline is identifying true mSTRs, we performed a more detailed analysis of mSTRs identified in the highly characterized NA12878 sample. To evaluate these mSTRs, we compared to an orthogonal dataset of haplotagged PacBio HiFi long reads (mean coverage *∼*30*×*) available for the same individual (**Methods**). Notably, although PacBio HiFi shows high accuracy at most regions, they have elevated error rates at homopolymers (Wenger *et al*., 2019), suggesting repeat counts obtained from PacBio reads at those loci may not serve as an accurate ground truth dataset. Additionally, we observed that inferred stutter error rates in short reads are highest at homopolymer STRs (**Supplementary Fig. 7**). Therefore, results below are reported separately for non-homopolymer vs. homopolymer STRs.

We reasoned that true mosaic alleles with sufficiently high variant allele fractions should be observed in both datasets, and that the mosaic allele should typically only occur on long reads from one of the two haplotypes at a locus (**Supplementary Fig. 14**). On the other hand, inferred mosaic alleles that are actually due to stutter or other error sources might be found on both haplotypes. After filtering, our analysis above had identified 440 candidate autosomal mSTRs, 400 (91%) of which occur at homopolymer loci. Of these, we deemed 16 (119) corresponding to 40% (27%) of candidate non-homopolymer (homopolymer) mSTRs to have sufficient PacBio HiFi coverage (at least 10 reads per haplotype) to attempt validation.

For each candidate mSTR, we examined the percentage of long reads from each haplotype supporting the inferred mosaic allele (*C*) (**Fig. 3A-B**) and classified calls into three categories. Category I, corresponding to 75% (50%) of non-homopolymers (homopolymers), consists of calls for which *C* is only identified in HiFi reads from a single haplotype, representing potential true positives. For these mSTRs, variant allele fractions estimated from short reads are strongly correlated with those observed in the HiFi reads (Pearson r=0.76, two-sided P=0.0040 for non-homopolymers and r=0.51, P=3.44e-05 for homopolymers; **Fig. 3C-D**). Category II, corresponding to 12.5% (47%) of non-homopolymers (homopolymers), consists of calls for which *C* is supported by at least one HiFi read from each haplotype, representing potential false positive calls. Category III, corresponding to 12.5% (3%) of non-homopolymers (homopolymers), consists of calls for which *C* is not supported by long reads on either haplotype. This could indicate an incorrect mSTR call, but could also originate from insufficient coverage at mSTRs with low variant allele fractions.

**Fig. 3.**
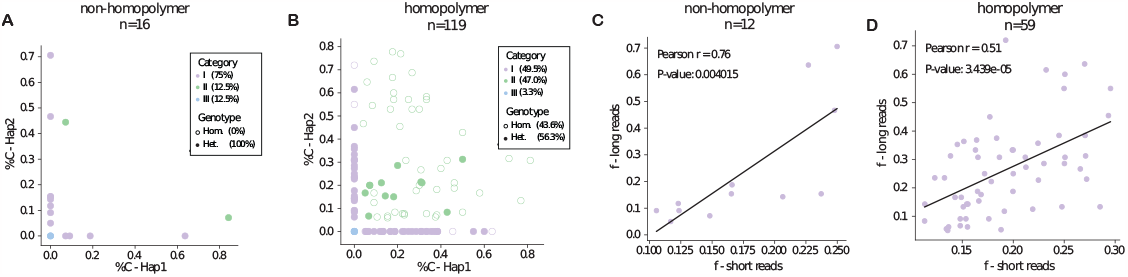
Validating candidate mSTRs identified in short reads from NA12878 (HG001) with PacBio HiFi reads. (A-B) Mosaic allele support on long reads from each haplotype. The x-axis and y-axis show the percentage of reads on each haplotype matching the mosaic allele for non-homopolymer (A) and homopolymer (B) loci. mSTRs were classified as Category I (purple; mosaic support on a single haplotype, potential true positives), Category II (green; mosaic support on both haplotypes, potential false positives), and Category III (blue; no mosaic support, undetermined). (C-D) Correlation of mosaic fraction between short and long reads. Comparisons of estimated allele fractions (for the mosaic allele identified in short reads) from short reads (x-axis) vs. PacBio reads (y-axis) are shown for non-homopolymer (C) and homopolymer (D) mSTRs. Black lines denote the best fit line. Only mSTRs in Category I are included.

We further examined read support on each haplotype at remaining candidate mSTRs (**Supplementary Fig. 15**). This revealed that the majority of homopolymer mSTRs identified as likely false positives occurred at loci for which the germline genotype was called as homozygous and long reads from both haplotypes supported multiple different alleles, suggesting reads at these loci are error prone. We additionally observed across all loci that the majority of validated high-confidence mSTRs occur at STRs for which the germline genotype is heterozygous. This is consistent with our simulation results, in which true mosaic alleles with high mosaic allele fraction occurring at homozygous sites are incorrectly genotyped as heterozygous and therefore systematically missing from our mSTR callset. On the other hand, those with low allele fraction are unlikely to be detected at genome-wide significance.

Finally, we evaluated whether the long read validation results above could be obtained by chance. In some cases an mSTR observed in short reads may appear to be confirmed by HiFi data but may be observed simply due to indel errors in long reads. To test this, we determined how often mSTRs inferred from other CEU samples would appear to validate in long reads from NA12878. For each sample, we computed the percent of mSTRs classified as Type 1 (potential true positives) and the significance (-log10 pvalue) of the correlation between the mosaic fraction observed in short vs. long reads, and compared those to metrics obtained for NA12878 (**Supplementary Fig. 16**). In all cases, mSTRs obtained from NA12878 showed the highest validation metrics. This analysis shows that although some mSTRs may appear to validate in long reads by chance, validation rates are substantially higher when the correct target sample is used, indicating many mSTRs identified by prancSTR are likely true positives. Overall, in combination with the population-wide analysis performed above, our results suggest mSTRs identified at heterozygous sites at moderate *f* values are robust, whereas accurate identification of mosaicism at homopolymers or for sites with either high or very low mosaic allele fractions remains challenging with short read data.

## Discussion

Here we presented prancSTR, a method for genome-wide detection of somatic mosaicism at STRs from high throughput sequencing datasets. prancSTR can accurately identify mSTRs without the need for a matched control sample. It has highest power to detect mSTRs with mosaic allele fractions of approximately 10-20% in PCR-free datasets with 30-50*×* coverage, but could detect reproducible mSTR sites with mosaic allele fractions as low as 7%. We applied prancSTR to identify mSTRs using PCR-free short read data for NA12878. Validation with orthogonal long read (Pacbio Hifi) data supported 75% and 50% of high-confidence mSTR calls at non-homopolymers and homopolymers, respectively, at sites with sufficient long read coverage. Application of prancSTR to population-scale short read WGS for the 1000 Genomes derived from lymphoblastoid cell lines identified hundreds of mSTRs per cell line with broadly consistent mSTR patterns across populations.

prancSTR is a versatile tool that can be used to detect mosaicism in a variety of settings, including PCR-free or PCR+ short read sequencing, as long as accurate stutter error parameters are available. It can also be applied as-is to PacBio Hifi datasets, which are becoming increasingly widely available and can be genotyped using HipSTR as done here. prancSTR as well as the read simulation method developed here (simTR) have been packaged into our existing toolkit, TRTools (Mousavi *et al*., 2021), enabling easy integration with other TR analysis tools. It is currently compatible with STR genotypes output by HipSTR (Willems *et al*., 2017), but could be easily modified to work downstream of other STR genotypers provided they output diploid genotypes and read support for each observed allele.

Application of prancSTR genome-wide to WGS from 460 cell lines revealed interesting patterns of mSTRs. Our results broadly suggest mSTRs identified from short reads at non-homopolymers and at sites with germline heterozygous genotypes are most reliable, whereas homopolymers remain particularly challenging. Overall, we found an average of 76 and 577 non-homopolymer and homopolymer mSTRs per cell line, corresponding to mutation rates of approximately 10^*−*4^ and 10^*−*3^ mutations per STR per sample. Intriguingly, we identified multiple cell lines from each population with outlier mutation counts, and found strong correlation between the number of mSTRs at homopolymers vs. non-homopolymers. This suggests some cell lines have higher rates of STR instability than others, and that these trends are present across a broad set of loci. Although we could not determine the source of this variation, we hypothesize it could be due to differences in cell line passage history, in which cell lines that have been maintained for more passages accumulate more somatic variation. It is also possible this variation arose from differences in NGS library preparation across samples. Alternatively, individual-level variation in mutation patterns could arise from germline factors such as mutations in DNA repair genes, which we have observed previously in mice (Maksimov *et al*., 2023). Profiling somatic variation in larger sample sizes is likely needed to identify similar effects in humans. prancSTR currently faces multiple limitations. First, it relies on an upstream genotyper (here, HipSTR) to provide accurate germline genotype calls as input. We identified several scenarios where germline genotype calls may be problematic. In cases where a mosaic allele is present at high frequency, it may be indistinguishable from a germline allele and incorrectly genotyped as heterozygous, causing mosaicism to be missed. Further, particularly at loci with high stutter error rates or low coverage, a truly heterozygous site may be incorrectly genotyped as homozygous, causing prancSTR to incorrectly identify the second germline allele as mosaicism. As a result, mSTRs identified at heterozygous sites are likely more reliable. We anticipate these challenges will be largely alleviated by haplotagged long reads, which will make distinguishing heterozygous vs. homozygous sites easier. Notably, an advantage of relying on an upstream genotyper is that it significantly reduces compute time required to identify mSTRs compared to directly analyzing raw NGS reads. Second, prancSTR currently focuses on identifying mSTRs with a single high frequency mosaic allele. While this is likely to capture mosaic events at shorter STRs, longer repeats such as the Huntington’s Disease locus where mosaicism is known to play a role in disease pathogenesis tend to show a broad range of mosaic allele lengths (Swami *et al*., 2009) and will require extensions to the current model to detect. Third, similar to mosaicism detection tools for other variant types, prancSTR is limited by the coverage of current datasets, which is insufficient to detect most mosaic events below 5% frequency.

Overall, prancSTR can serve as a valuable method to characterize somatic mosaicism at STRs in a range of settings, including in healthy individuals or in disease settings such as microsatellite instability in cancer or neurological diseases where mosaicism is known to play a key role. Profiling mosaicism at population-scale from the large number of existing WGS datasets may also give insight into inherited factors driving differences in mSTRs patterns across individuals. We envision future extensions of this framework can allow for directly incorporating phase information from haplotagged reads or quantifying mosaicism at highly unstable repeats such as long Huntington’s alleles, which will further improve our ability to characterize STR mosaicism and its role in human health.

## Acknowledgements

We thank Alon Goren, Vineet Bafna, and Eric Mendenhall for helpful discussions about the manuscript. We thank Arya Massarat and Jonathan Margoliash for help integrating prancSTR and simTR into TRTools.

