## Supplemental Materials for "Genome-wide detection of somatic mosaicism at short tandem repeats"

### Supplementary Material - prancSTR

#### Supplementary Figures

##### Supplementary Figure 1

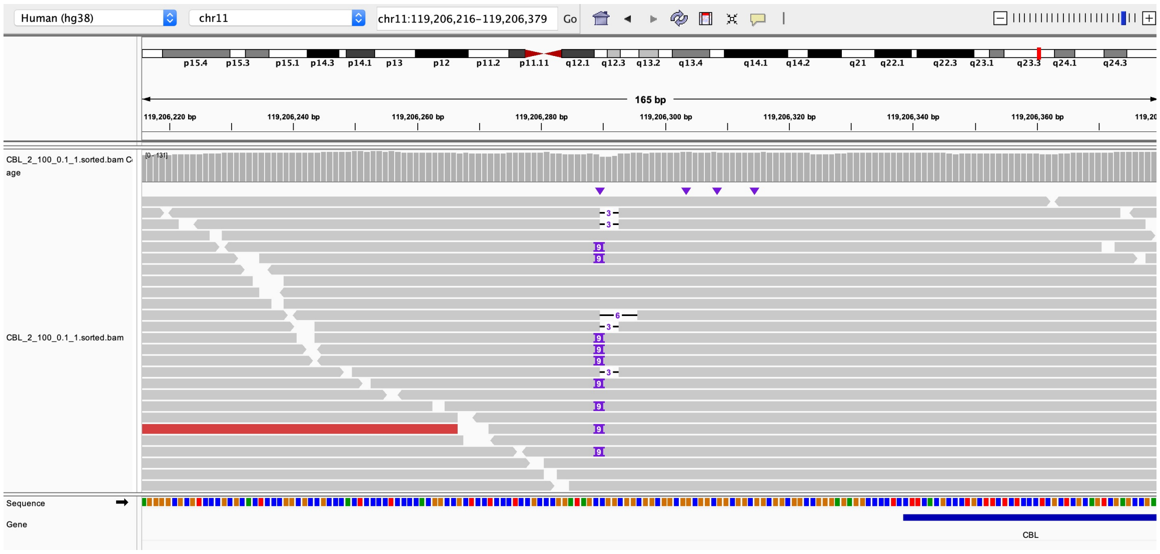


Figure 1: **Visualization of reads simulated by simTR.** Reads were simulated in a 1000bp window around a CGG repeat in *CBL* (hg38 chr11:119206289-119206322). The simulation was based on a germline genotype of 11 (reference allele) and 14 copies (reference + 9bp) of the repeat, and a mosaic allele with 10 copies (reference - 3bp) at 10% variant allele fraction. The visualization was produced using the Integrative Genomics Viewer (https://software.broadinstitute.org/software/igv/).

#### Supplementary Figure 2

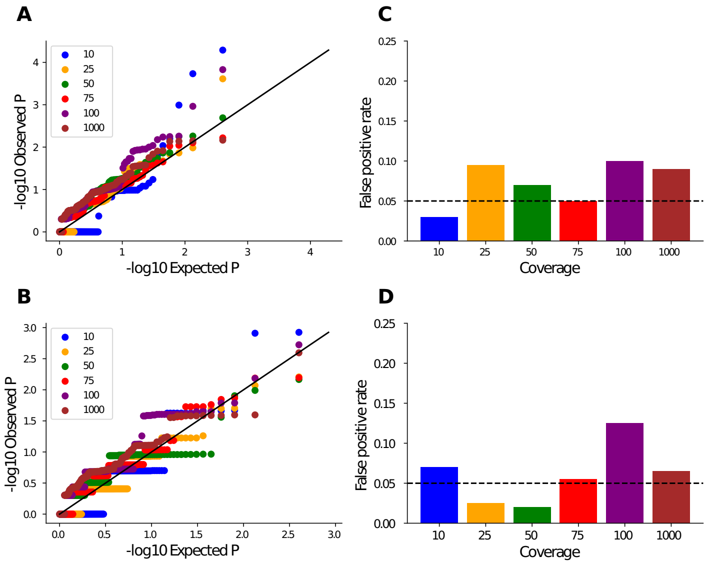


Figure 2: **prancSTR p-values are well-calibrated.** (A-B) represent the quantile-quantile (QQ) plots showing the distribution of p-values in simulations where the germline genotype is either heterozygous (A) or homozygous (B) under different coverage levels (denoted by dot colors) and where no mosaicism was simulated ($f = 0$). As expected, p-values follow the uniform distribution. (C-D) show the false positive rate, computed as the percent of null ($f = 0$) simulations (n=200) in which a significant p-value ($p<0.05$) was obtained. Bar color denotes coverage. Panels here are based on simulated read vectors.

#### Supplementary Figure 3

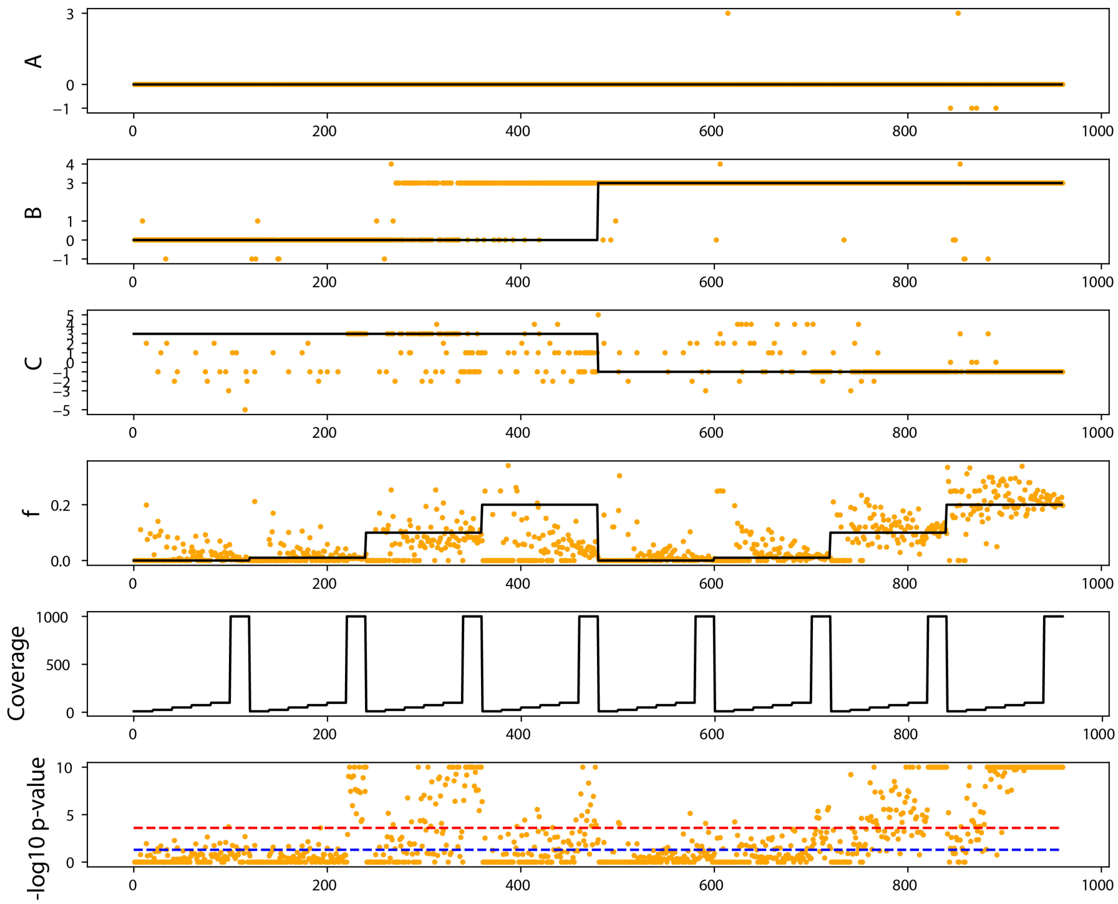


Figure 3: **Benchmarking prancSTR using simulated reads - CBL locus**. We used simTRto simulate reads which were input to HipSTR followed by prancSTR to detect STR mosaicism under a range of settings. Results are shown for a CGG repeat near *CBL* (hg38 chr11:119206289-119206322). We varied the germline genotypes ($A$ and $B$; top two panels), mosaic allele ($C$; third panel), mosaic allele fraction ($f$; fourth panel), and target coverage (fifth panel). In each of the top five panels, black lines show the simulated value. Each simulation setting was performed 20 times. Orange dots denote estimated parameter values. Germline genotypes ($A$ and $B$) are those estimated by HipSTR. Estimated values of $f$ and $C$ are obtained from prancSTR. The bottom plot shows the -log10 p-value obtained in each case by prancSTR, which tests the null hypothesis that $f = 0$. The blue dashed line denotes p=0.05. The red dashed line denotes p=0.00025, which is the genome-wide significance threshold used to identify mSTRs at FDR 5% when running on a single sample (NA12878).

**Supplementary Figure 4**

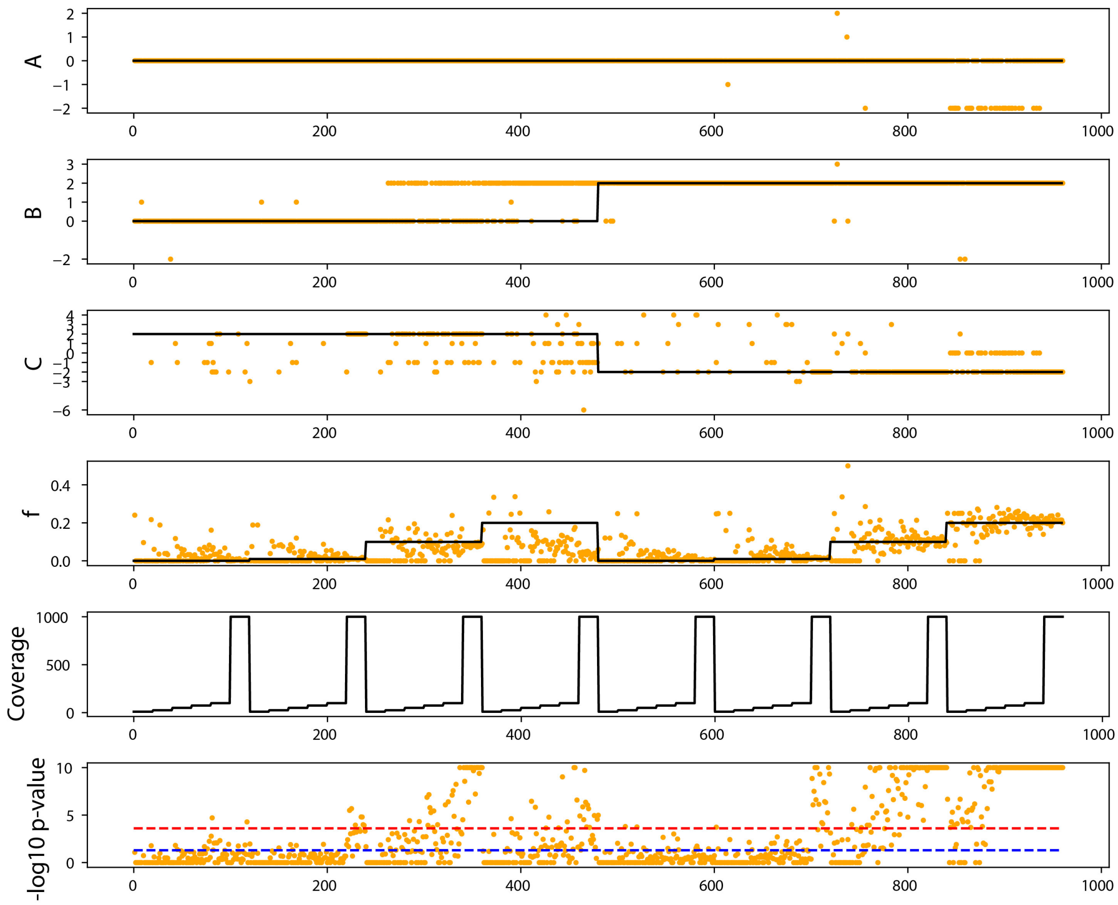


Figure 4: **Benchmarking using simulated reads - CSF1PO locus** Panels are same as in **Supplementary Fig. 3** except reads are simulated for the tetranucleotide (AGAT) CSF1PO locus (hg38 chr5:150076324-150076375).

#### Supplementary Figure 5

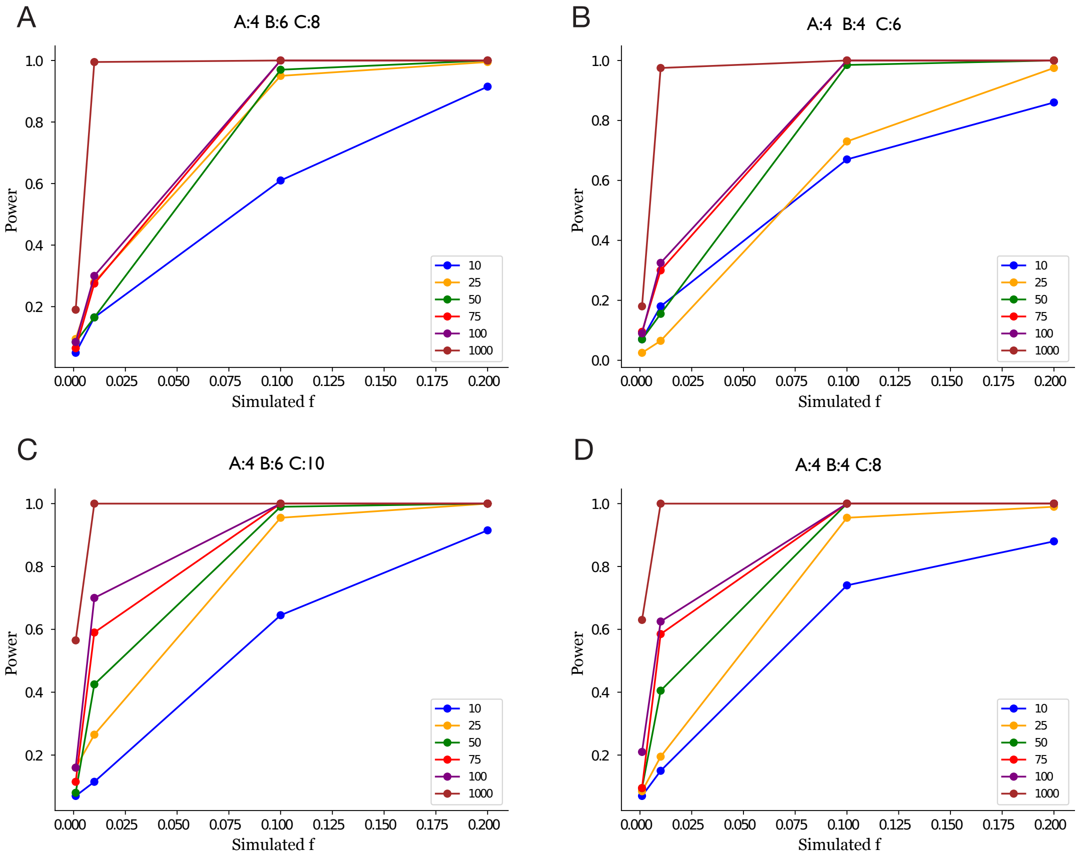


Figure 5: **Power of detection across different values of C**. (A-B) show the power of detection in a germline heterozygous (A) vs homozygous (B) case, where the mosaic allele $C$ is two repeat units away from the nearest germline allele. (C-D) show the power of detection in a heterozygous (C) vs homozygous (D) case, where the mosaic allele $C$ is four repeat units away from nearest germline allele. As in **Fig. 1D-E**, power is computed as the percent of simulations for which $p < 0.05$. Lines denote different coverage levels, where coverage gives the total number of reads spanning the STR of interest. Panels here are based on simulated read vectors.

#### Supplementary Figure 6

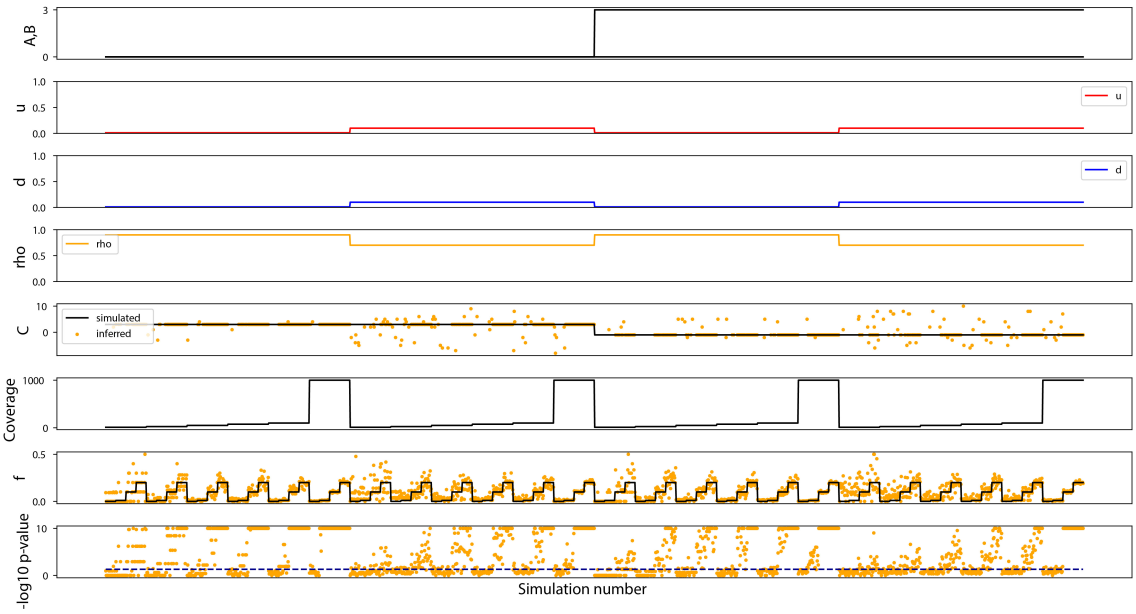


Figure 6: **Evaluating mSTR detection under additional simulation settings**. The top seven panels show simulated values for the germline genotype ($A, B$; top panel), stutter expansion probability ($u$; second panel), stutter contraction probability ($d$; third panel), stutter step size (rho; fourth panel), mosaic allele ($C$; fifth panel), coverage (sixth panel), and mosaic allele fraction ($f$; seventh penal). In each of the top seven panels, the lines shows the simulated value. Each simulation setting was performed 20 times. Orange dots denote estimated parameter values. The bottom plot shows the -log10 p-value obtained in each case by prancSTR, which tests the null hypothesis that $f = 0$. The dark blue dashed line denotes p=0.05. Panels here are based on simulated read vectors.

#### Supplementary Figure 7

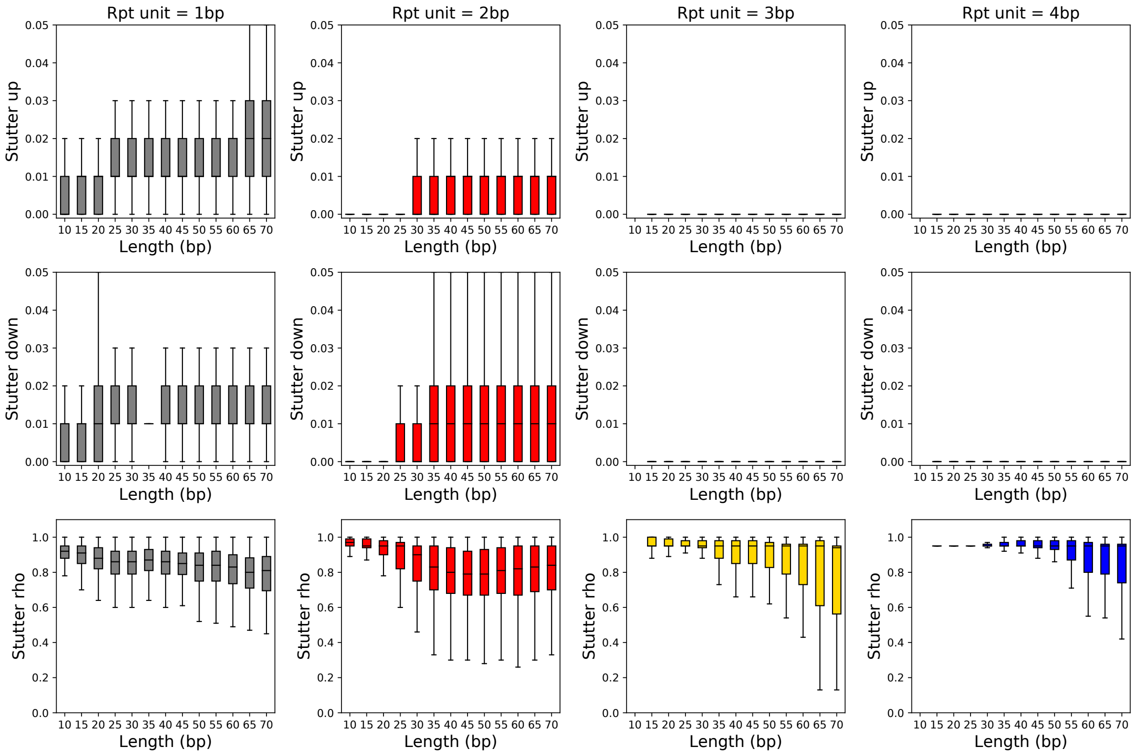


Figure 7: **Characterizing stutter parameters in the 1000 Genomes**. Per-locus stutter parameters were previously estimated from PCR-free high-coverage WGS for 1000 Genomes samples using HipSTR (Ziaei Jam *et al.*, 2023). Boxplots in each row show the distribution of the three stutter parameters output by HipSTR (top=INFRAME_UP, middle=INFRAME_DOWN; bottom=INFRAME_PGEOM) for STRs with different repeat unit lengths (columns: gray=homopolymers; red=dinucleotides; gold=trinucleotides; blue=tetranucleotides). Horizontal lines show median values, boxes span from the 25th percentile (Q1) to the 75th percentile (Q3). Whiskers extend to Q1–1.5*IQR (bottom) and Q3+1.5*IQR (top), where IQR gives the interquartile range (Q3-Q1).

#### Supplementary Figure 8

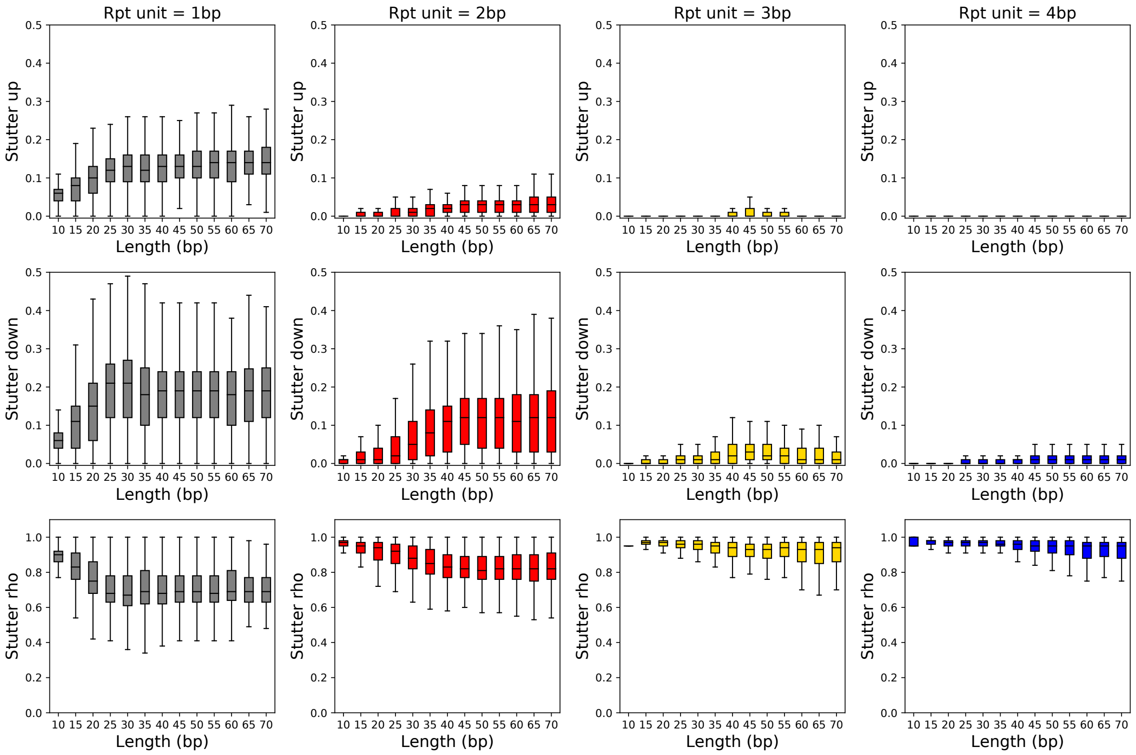


Figure 8: **Characterizing stutter parameters in the H3Africa**. Per-locus stutter parameters were previously estimated from PCR+ high-coverage WGS for individuals from the H3Africa cohort using HipSTR (Ziaei Jam *et al.*, 2023). Panels showing distributions of stutter parameters are the same as in **Supplementary Fig. 7** except based on H3Africa data.

**Supplementary Figure 9**

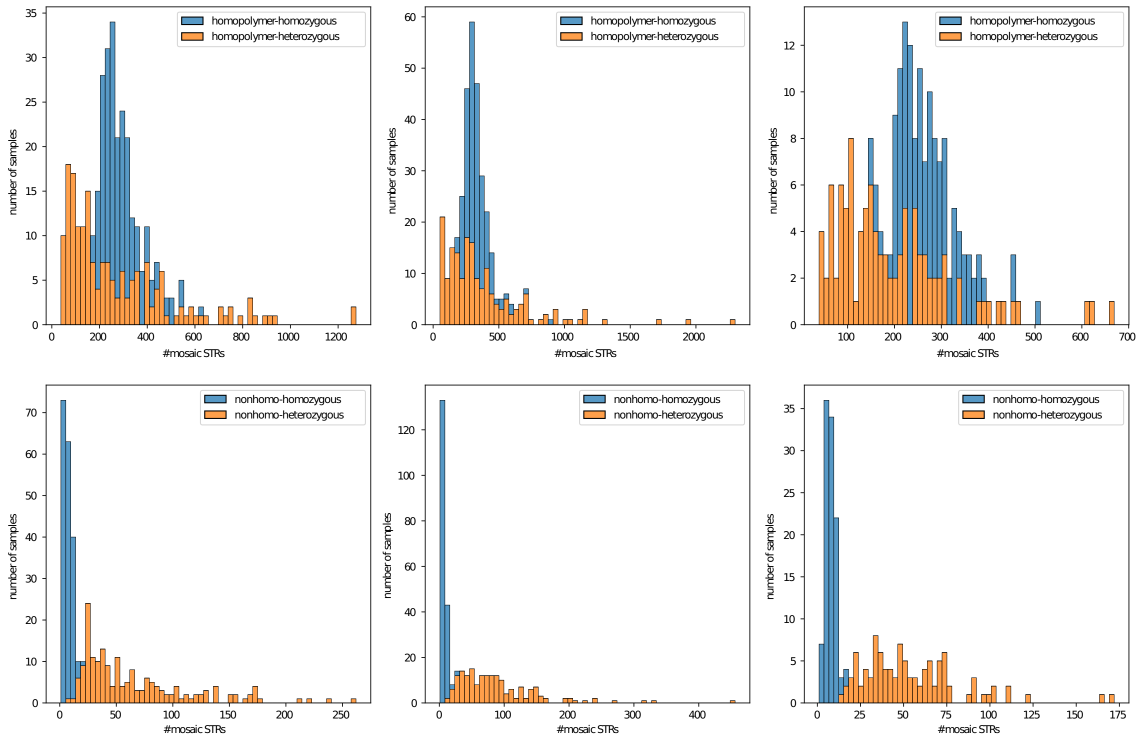


Figure 9: **Distribution of mSTRs detected per sample in each population** Histograms show the number of mSTRs detected per cell line after quality filtering (**Methods**). Blue=mSTRs at germline homozygous sites and orange=mSTRs at germline heterozygous sites. Top and bottom plots show counts at homopolymer and non-homopolymer STRs, respectively. Data is shown separately for CEU (left), YRI (middle), and CHB (right).

#### Supplementary Figure 10

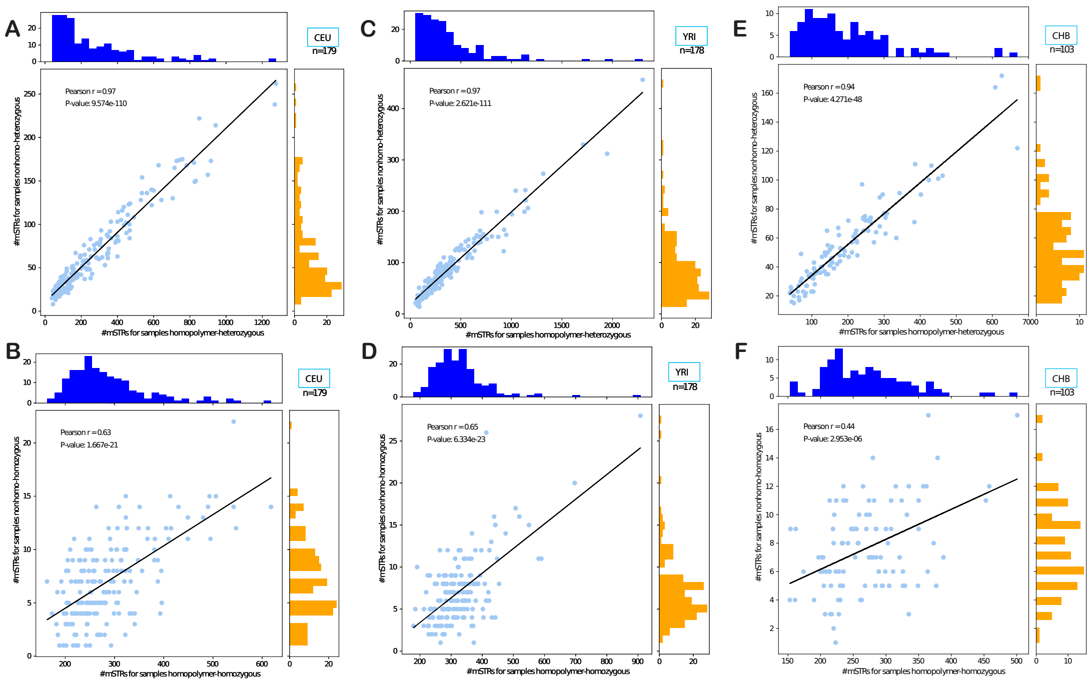


Figure 10: **Distribution of mSTRs detected per sample for homopolymers vs non-homopolymers**. Scatter plots show the number of mSTRs detected per cell line after quality filtering (**Methods**) in homopolymers (x-axis) vs non-homopolymers (y-axis). Top and bottom plots are restricted to mSTRs occurring at germline heterozygous and homozygous sites, respectively. Data is shown separately for CEU (left), YRI (middle), and CHB (right). Black lines show the best linear fit. Pearson correlation coefficients and corresponding two-sided p-values are annotated in each plot.

#### Supplementary Figure 11

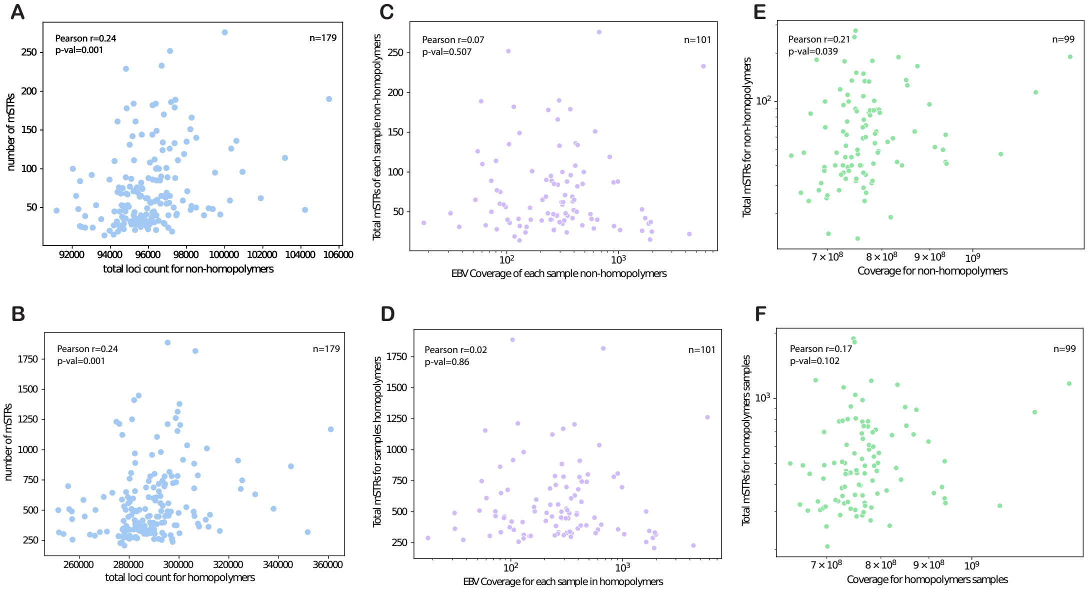


Figure 11: **Evaluation of factors influencing the number of mSTRs per cell line**. Scatter plots show the relationship between various factors (x-axis) and the number of mSTRs detected after filtering (y-axis). A-B=total number of loci analyzed per cell line, C-D=EBV coverage, E-F=sequencing coverage. Top and bottom plots are restricted to mSTRs occurring at non-homopolymers and homopolymers, respectively. Data is shown for CEU only. Other populations showed similar trends (not shown). Pearson correlation coefficients and corresponding two-sided p-values are annotated in each plot.

#### Supplementary Figure 12

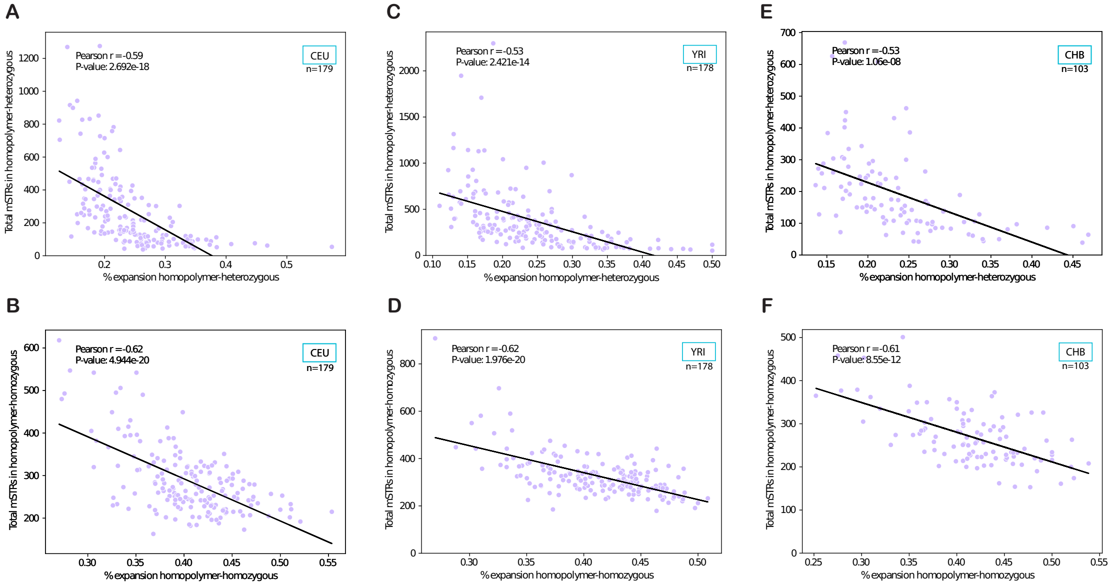


Figure 12: **Relationship between expansion bias and number of mSTRs per cell line.** Scatter plots show the % of mSTRs that are expansions (x-axis) vs. the total number of mSTRs detected per cell line (y-axis). Top and bottom plots are restricted to mSTRs occurring at germline heterozygous and homozygous sites, respectively. Data is shown separately for CEU (left), YRI (middle), and CHB (right). Black lines show the best linear fit. Pearson correlation coefficients and corresponding two-sided p-values are annotated in each plot. Data is only shown for homopolymers.

#### Supplementary Figure 13

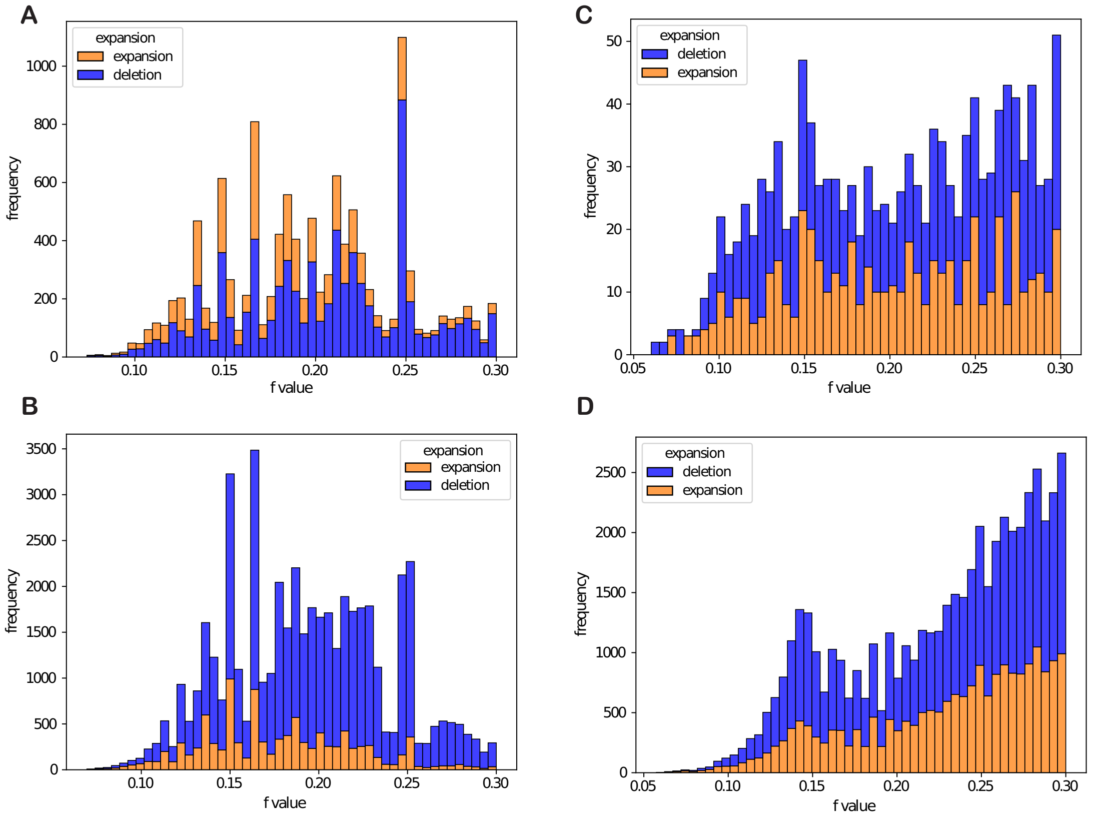


Figure 13: **Distribution of f across all mSTRs detected in 1000Genomes cell lines.** Top and bottom plots are restricted to mSTRs occurring at non-homopolymers and homopolymers respectively. Left and right plots represent mSTRs at germline heterozygous and homozygous sites, respectively. The histograms are further broken down by expansions (orange) and deletions (blue). Bars are stacked, such that the y-axis value of each bar denotes the total number of mSTRs falling in each bin of f-values. Data is shown for CEU only. Other populations showed similar trends (not shown).

**Supplementary Figure 14**

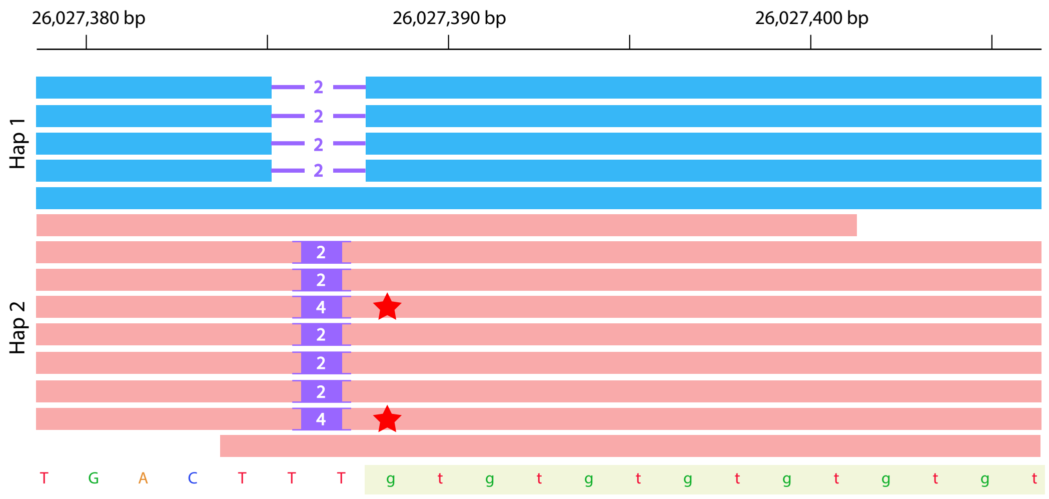


Figure 14: **Schematic representation of mosaicism validation with long reads.** PacBio HiFi reads are haplotagged as belonging to either of the two haplotypes in a sample. The highlighted region denotes the STR region. Bars indicate deletions and purple rectangles indicate insertions compared to the reference genome. Mosaic alleles (red star) are typically expected to occur on only one of the two haplotypes.

**Supplementary Figure 15**

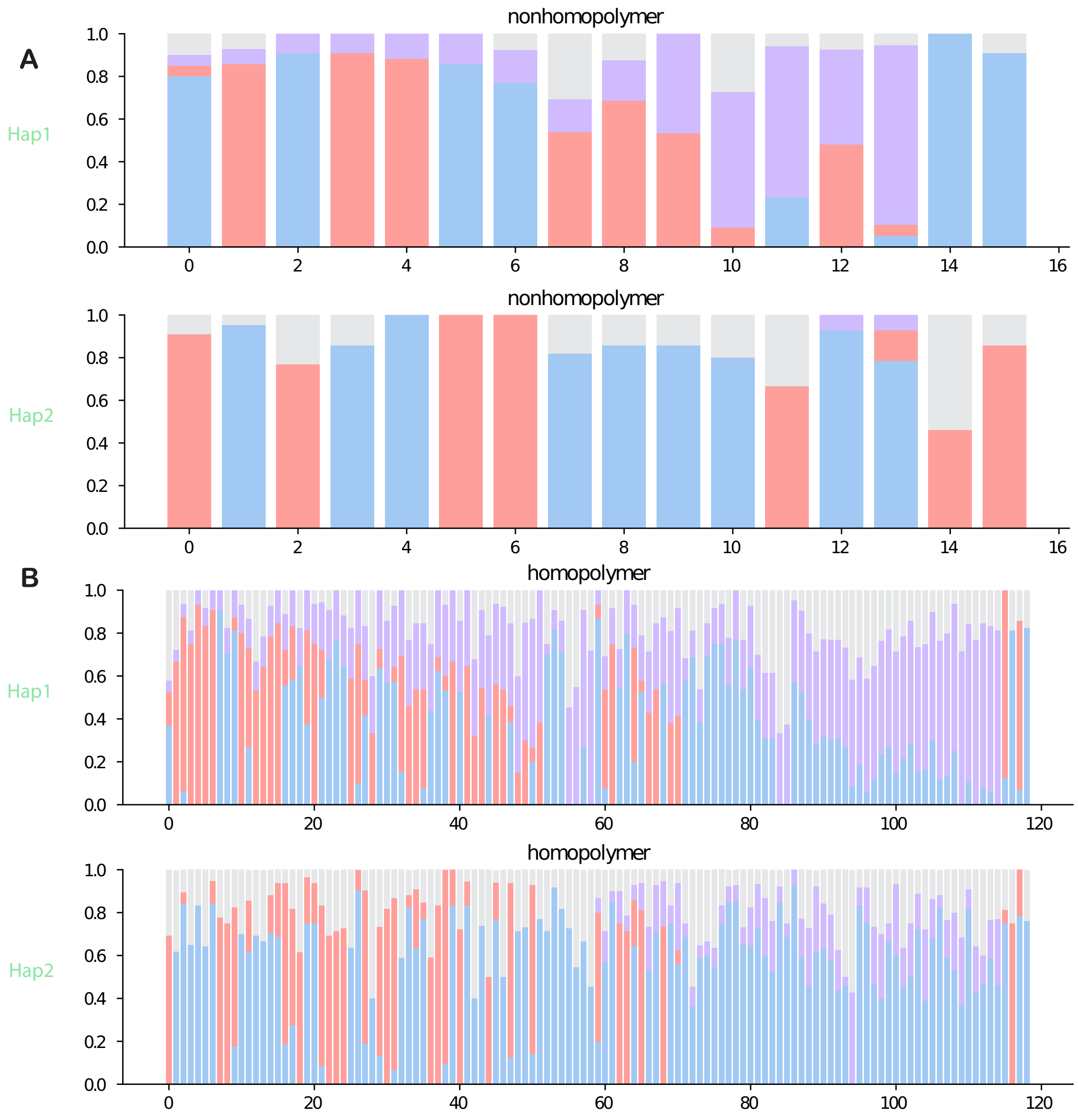


Figure 15: **Inspection of per-locus long read support for candidate mSTRs inferred from short reads in NA12878.** Each bar shows the percentage of PacBio Hifi reads on each haplotype supporting the germline genotype alleles (scarlet and blue), mosaic allele (lavender), or other alleles (gray). (A) shows the two haplotypes at each candidate non-homopolymer mSTR and (B) show the two haplotypes at homopolymer mSTRs. The order of the two haplotypes at each locus is arbitrary.

**Supplementary Figure 16**

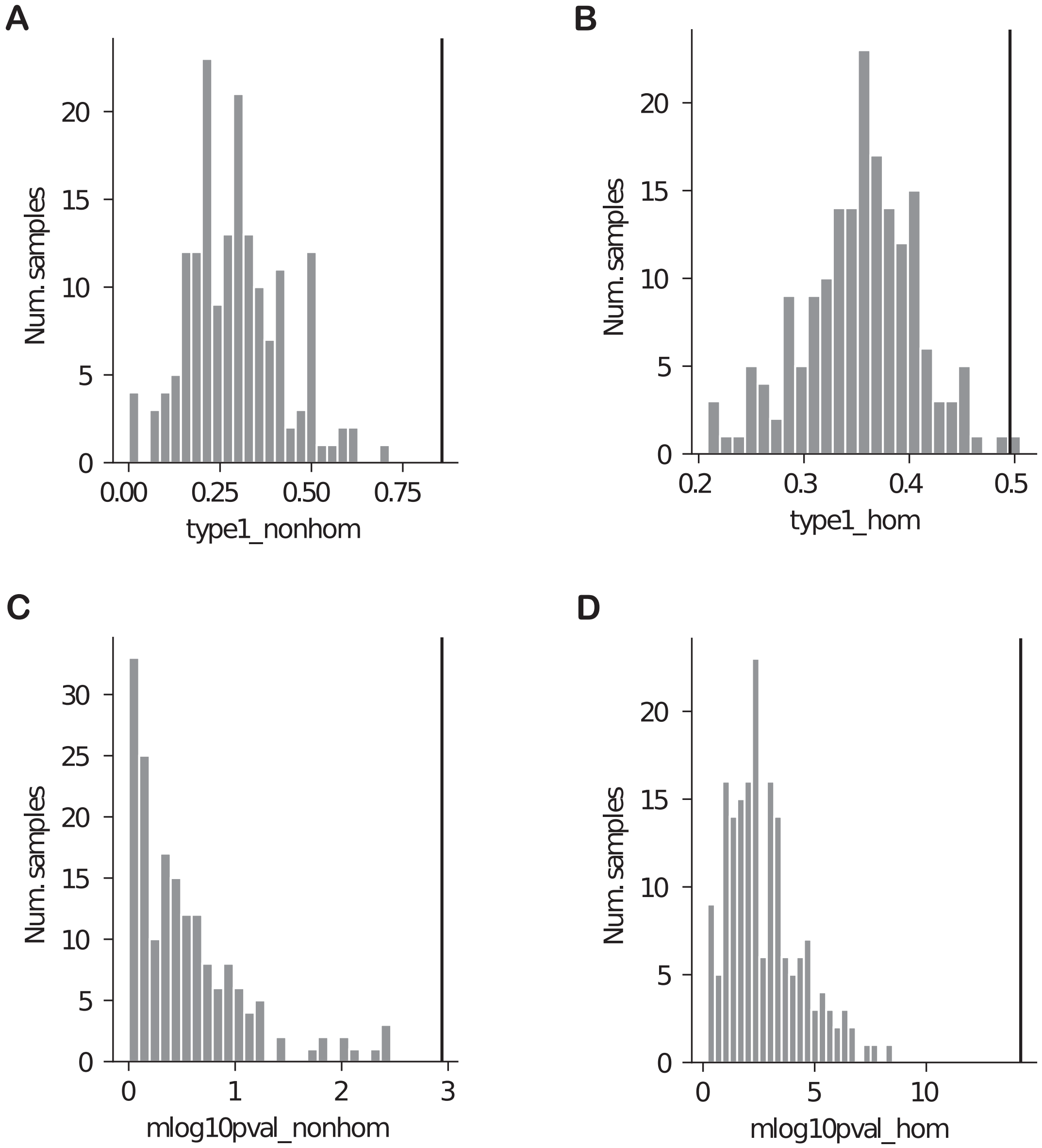


Figure 16: **Comparison of NA12878 long read validation metrics in other CEU samples.** As a negative control, we performed long read validation analysis using mSTRs identified in other CEU samples from short reads and compared metrics to those obtained from the matching sample (NA12878). Histograms (gray bars) show the distribution of each metric in all CEU samples except NA12878. The black line indicates the metric computed in NA12878. (A) and (B) show the percentage of short read mSTRs identified as Type 1 (potential true positives) in long reads for non-homopolymer and homopolymer loci, respectively. (C) and (D) show the -log10 p-value of the Pearson correlation between the mosaic allele fraction estimated from short reads vs. that observed in long reads for non-homopolymer and homopolymer loci, respectively.
